# CellProfiler 4: Improvements in Speed, Utility and Usability

**DOI:** 10.1101/2021.06.30.450416

**Authors:** David R. Stirling, Madison J. Swain-Bowden, Alice M. Lucas, Anne E. Carpenter, Beth A. Cimini, Allen Goodman

## Abstract

**Background:** Imaging data contains a substantial amount of information which can be difficult to evaluate by eye. With the expansion of high throughput microscopy methodologies producing increasingly large datasets, automated and objective analysis of the resulting images is essential to effectively extract biological information from this data. CellProfiler is a free, open source image analysis program which enables researchers to generate modular pipelines with which to process microscopy images into interpretable measurements.

**Results:** Herein we describe CellProfiler 4, a new version of this software with expanded functionality. Based on user feedback, we have made several user interface refinements to improve the usability of the software. We introduced new modules to expand the capabilities of the software. We also evaluated performance and made targeted optimizations to reduce the time and cost associated with running common large-scale analysis pipelines.

**Conclusions:** CellProfiler 4 provides significantly improved performance in complex workflows compared to previous versions. This release will ensure that researchers will have continued access to CellProfiler’s powerful computational tools in the coming years.

## Background

Microscopy can be used to capture images which contain a wealth of information that can inform biomedical research. Image analysis software can allow scientists to obtain quantitative measurements from images that are otherwise difficult to capture via subjective observation. The increasing use of automated microscopy now allows researchers to capture images of samples treated with many thousands of individual compounds or genetic perturbations. Scientists increasingly image cells in 3D or across time series; this expanding bulk of raw data necessitates automated processing and analysis. Such analysis is best achieved through using software to perform automated detection of cells or organisms and extract quantitative metrics which objectively describe the specimens.

Many microscopes are now sold with accompanying proprietary analysis packages, such as MetaMorph (Molecular Devices), Elements (Nikon), Zen (Zeiss) and Harmony (Perkin Elmer). These ecosystems are powerful but can lack the flexibility to work with data from other manufacturers’ equipment. Cost of these proprietary solutions can also limit accessibility, and their closed-source nature can obscure exactly how scientists’ data is being analyzed. Free, open-source software packa(1)ges such as ImageJ, CellProfiler, QuPath, Ilastik and many others have therefore become popular analysis tools used by researchers. ImageJ is the most widely-used package and excels in performing analysis of single images, assisted by a vast array of community-developed plugins (1). Numerous smaller packages are tooled towards specific types of data: for example, QuPath is a popular program geared specifically towards pathology applications (2), while Ilastik delivers an interactive machine learning framework to assist users in segmenting images (3).

In 2005 we introduced CellProfiler, an open-source image analysis program which allows users without specific training to automate their image analysis by using modular processing pipelines (4). CellProfiler has been widely adopted by the community, and is currently referenced more than 2000 times per year. Built-in modules provide a diverse array of algorithms for analyzing images, which can be further extended through the use of community-developed plugins. In an independent analysis of 15 free image analysis tools CellProfiler scored highly in both usability and functionality (5). Our previous release, CellProfiler 3, introduced support for analysis of 3D images to further expand the tool’s applications (6). However, some popular features from CellProfiler 2 could not be brought forward into that release and certain modules struggled to operate efficiently in 3D pipelines.

## Implementation

CellProfiler was originally written in MATLAB, but in 2010 was rewritten in Python 2, which reached its official end-of-life in 2020. In order to ensure ongoing compatibility with future operating systems we ported the software to the Python 3 language to create CellProfiler 4. This provided the opportunity for a broader restructuring of the software’s code to improve performance, reliability and utility. CellProfiler 4 is available for download at cellprofiler.org.

As part of the migration to Python 3, we split the CellProfiler source code into two packages: cellprofiler and cellprofiler-core. The new cellprofiler-core package contains all the critical functionality needed to execute CellProfiler pipelines, whereas the cellprofiler repository now primarily contains the user interface code and built-in modules. The core package has been developed to introduce a stable API which will allow users to access CellProfiler’s functionality as a Python package within popular environments such as Jupyter (7) and for future integration with other packages and software suites.

### User interface refinements

Guided by feedback from biologists, we have made several improvements to the CellProfiler user interface with the goal of making the software more accessible and easier to use. The basic 3D viewer introduced in CellProfiler 3.0 has now been replaced with a more fully-featured viewer which allows users to inspect any plane in a volume (Figure 1A). We have also expanded the figure contrast dialogs to give users more granular control over how images are displayed in both 2D and 3D mode (Figure 1B). These changes will help users to better visualize and understand their data.

**Figure 1.**
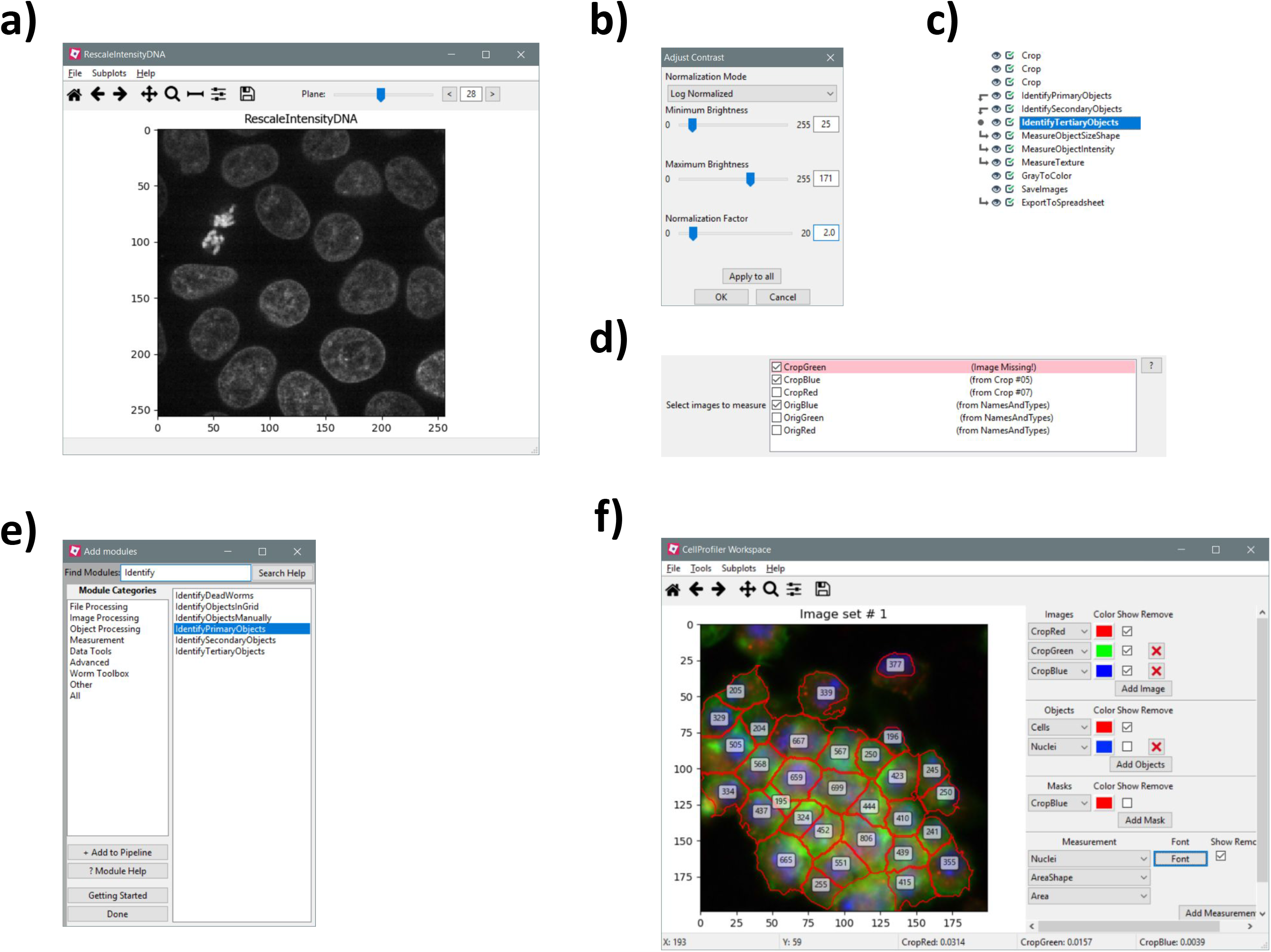
User interface refinements in CellProfiler 4. (A) The new 3D viewer window with plane controls in the top right. (B) Contrast and normalization adjustment popup available with any image window. (C) Interface displayed when the “trace” command is called on a module. Arrow icons on the left represent modules which provide data to or use data from the selected module (dot icon). (D) Selection widget for choosing multiple images for analysis. Images sourced from disabled or missing modules are highlighted. (E) “Add modules” pane in search mode, the modules list is filtered based on the text entered into the search box. (F) The Workspace Viewer module displaying a custom overlay of data from the example pipeline.

Other changes make it easier to develop and configure pipelines. We added an interface to visualize which modules produce inputs needed by, or use outputs from, a module of interest, which will aid in modifying complex pipelines (Figure 1C). We also revised the interface for selecting multiple images for analysis within a module, replacing dropdown menus with a checklist in which multiple images can be selected quickly and efficiently (Figure 1D). Furthermore, a new search filter in the “Add module” popup allows users to more easily find desired modules by module name rather than by category (Figure 1E).

We also restored some features which were previously lost in the migration from CellProfiler 2 to CellProfiler 3. Most notably, we rebuilt the Workspace Viewer, where users construct a customized view of their data and can stay focused on a specific region of interest as the pipeline is modified (Figure 1F), making it much simpler to monitor and refine segmentation of problematic regions of an image. In addition, new icons in the Test Mode pipeline interface provide a stronger visual indication of which module is currently about to be executed, and provide the ability to return to and execute earlier modules in the pipeline. This replicates and replaces the functionality of the slider widget from CellProfiler 2, which could not be carried forward into CellProfiler 3 but was popular with users.

### New and restored features

In CellProfiler 4 we introduced several new analysis features and settings. A common workflow issue we identified was that analysts often segment highly variable objects in multiple stages (such as segmenting and masking out bright objects to aid segmentation of similar-but-dimmer objects), but previous versions could not simply treat resulting segmentations as a single object set when performing and exporting measurements. To resolve this we added the CombineObjects module to allow users to merge sets of objects which have been defined separately. A key issue when designing this module was how to handle objects that would overlap if the sets were merged, therefore we built several strategies detailed in Figure 2. The resulting merged set can then be carried forward throughout the pipeline without the need to merge measurement tables outside of CellProfiler.

**Figure 2.**
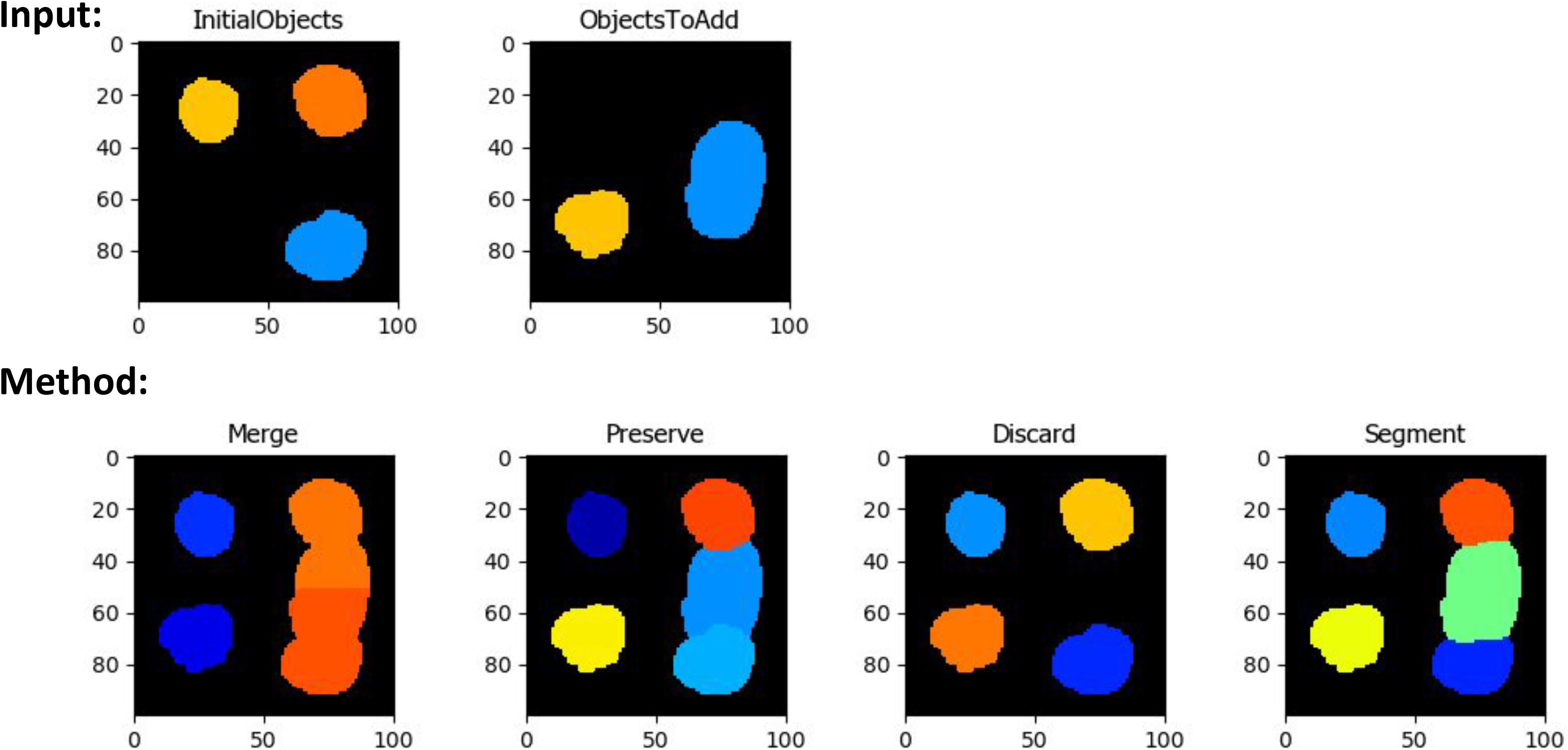
Approaches for combining object sets within the CombineObjects module. Results represent the output from different methods available within the module. “Merge” will join touching objects and distribute conflicting regions to the nearest object from the initial set. “Preserve” will add only regions of objects from the second set which did not overlap with the initial set. “Discard” will only add objects with no overlap. “Segment” will add both object sets and re-segment disputed regions.

Many users were disappointed with the loss of the RunImageJ module (8) in CellProfiler 2.2; we have now replaced it with the new RunImageJMacro module. The new module allows a user to export images from CellProfiler into a temporary directory, execute a custom ImageJ macro on that directory and then automatically import resulting processed images back into CellProfiler. In practice this will allow users to access ImageJ functions and plugins within a CellProfiler pipeline, greatly expanding its interoperability. Unlike its predecessor, the RunImageJMacro module relies on the user’s copy of ImageJ rather than a built-in copy. This allows users to take advantage of any new ImageJ upgrades and simultaneously poses less danger to CellProfiler’s stability because releases between the two softwares need not be kept in sync.

We also upgraded several existing modules. We rewrote the Threshold module to allow all pre-existing threshold strategies to be used in ‘adaptive’ mode, giving users more options in images with highly-variable background. We have also added the Sauvola local thresholding method as an alternative adaptive strategy (9). Previous versions of CellProfiler 2 shipped a version of the Otsu thresholding method that log-transformed the data before applying the threshold; this assisted in the thresholding of dim images, but led users to question why our Otsu values did not match those from other libraries such as scikit-image (10). This inconsistent behavior could be confusing to users, so we began the process of updating that implementation in CellProfiler 3 and completed it in CellProfiler 4. We added a dedicated setting to log transform image data during application of any thresholding method. These new options will assist users in segmenting challenging images.

### New Measurements

We overhauled some measurement modules in CellProfiler 4. We redesigned MeasureObjectSizeShape to record additional measurements now available in scikit-image, including bounding box locations, image moments and inertia tensors, producing up to 60 new shape measurements per object. We anticipate that these new features may be of particular value for training machine learning models, which play an increasingly important role in performing object classification on large data sets. In addition to new features, several of the previously 2D-exclusive measurements, such as Euler Number and Solidity, are now also available when working with 3D images. Together these expanded measurements provide researchers with even more metrics with which to investigate cellular phenotypes.

## Results

### Performance improvements

A key focus in producing CellProfiler 4 has been improving performance of the software and addressing common issues encountered by users. We revised our build packaging process to more reliably bundle CellProfiler’s Java dependencies so that additional software and system configuration is no longer required to use the program. In doing so we also optimized the program’s startup sequence, which provided a substantial improvement in the time taken to initialize the software (Figure 3A). Another critical area of focus for improvement has been in file loading (input/output, or I/O operations). Combined improvements in Python’s underlying directory scanning functions and optimizations to CellProfiler’s image loading procedures have dramatically reduced the time needed to add large folders of images to the file list. This is particularly noticeable when using networked storage.

**Figure 3.**
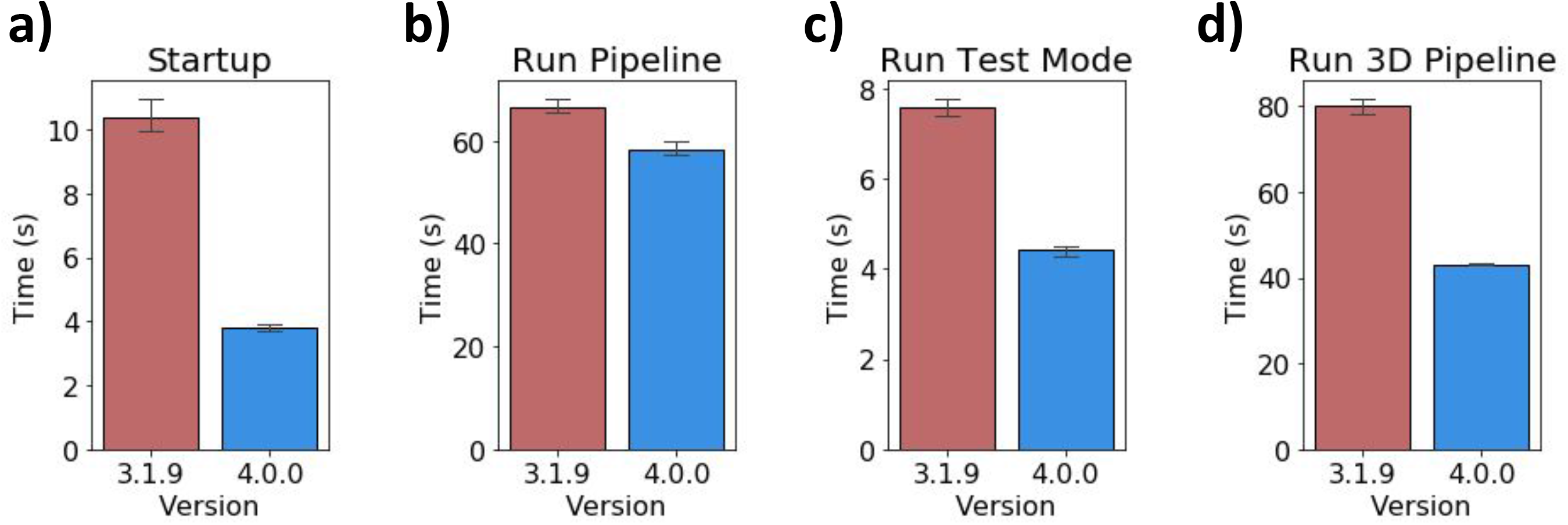
General performance in CellProfiler 3 vs CellProfiler 4. Results represent independent runs on a machine running Windows 10, using 1 worker process. (A) Time from launching the CellProfiler executable to display of the full GUI (n=5). (B) Time taken to run the ExampleFly pipeline in Analysis Mode (n=3). (C) Time to run the ExampleFly pipeline in Test Mode (n=5). (D) Time to run the 3D monolayer tutorial pipeline in Analysis Mode (n=3).

In our performance testing of an example analysis pipeline, overall performance was similar to CellProfiler 3 (Figure 3B). However, executing this pipeline in Test Mode was inhibited by unnecessary user interface updates between running individual modules. Optimizing the UI updates sent during test mode reduced the time taken to run an image set in this mode (Figure 3C).

Running more complex analysis workflows such as 3D segmentation and the commonly used Cell Painting assay (11) was time-consuming in CellProfiler 3. We therefore aimed to identify and refine modules which displayed long execution times in these scenarios.

Optimizations across all modules produced a 50% performance improvement when running 3D pipelines such as the 3D monolayer tutorial dataset (Figure 3D) (12). Within 3D workflows we had identified the MedianFilter module as being particularly slow to process. By switching to the new scipy.ndimage filter implementation we were able to substantially reduce the time taken to process each image (Figure 4A).

**Figure 4.**
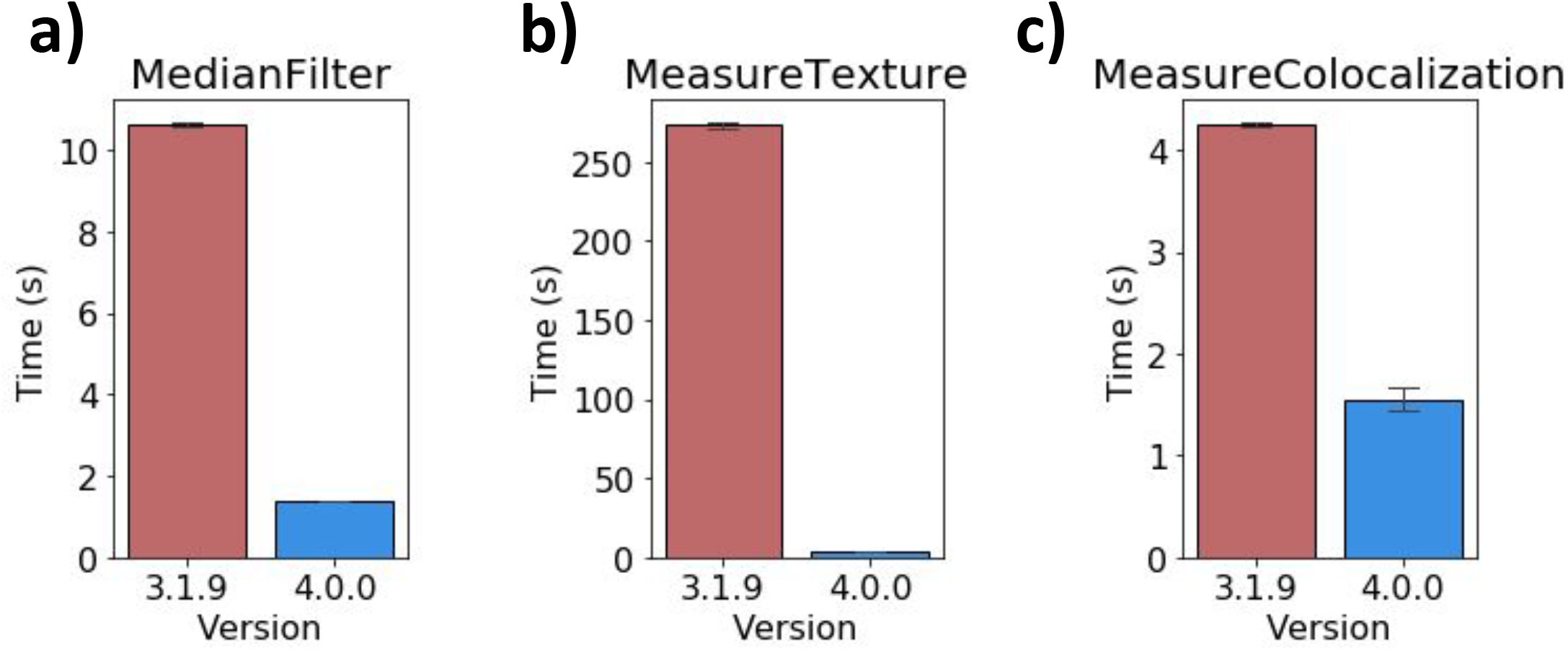
Module-specific performance improvements. Results from individual module testing on a machine running Windows 10. (A) Execution time for the MedianFilter module running within the 3D Monolayer pipeline (n=5). (B) Execution time when running per-object texture measurements on data from the ExampleFly pipeline (n=5). (C) Execution time when running MeasureColocalization on 8-bit images from the ExampleFly pipeline (n=5).

Another key target was the MeasureTexture module, which exhibited long run times when performing per-object measurements. Analysis revealed that this was caused by per-object functions processing full-size masked arrays for each object to be measured. To improve performance we adjusted these functions to produce and process arrays cropped down to the particular region of interest for each object. In our testing this reduced the time taken to analyze each image from minutes down to seconds, without any change in the resulting measurements (Figure 4B).

Major gains were also made in measurement of the Costes Colocalization Coefficient in the MeasureColocalization module. This statistic requires the calculation of Costes’ automatic threshold, which is generated by thresholding the two images to be compared and then serially reducing the thresholding value until the Pearson R correlation between the two thresholded images drops below a value of 0. Our original implementation would reduce the candidate value in images scaled 0-1 by 1/255 at each step, which was inappropriate for 16-bit images containing 65,536 grey levels rather than the 256 present in 8 bit images. Testing 65,536 candidate thresholds in 16-bit images would be excessively slow, so we introduced optional alternative implementations of the Costes automated thresholding method to resolve this inefficiency. Our first optimization maintained the canonical strategy of evaluating every possible threshold, but only measured the Pearson R correlation of the thresholded images if the new value produced a different total number of thresholded pixels than the previous value. We termed this “accurate” mode, but in images with large numbers of unique pixel values performance was unacceptably slow. We therefore introduced “fast” mode to the module, in which the candidate threshold is decreased in larger steps if the previous Pearson R value was substantially higher than 0. This improved performance when working with 8-bit images (Figure 5A), but was still inefficient with 16-bit images (Figure 5B). We subsequently devised an alternative implementation, dubbed “faster” mode, in which a weighted bisection search algorithm is used to consecutively narrow a window of possible target thresholds. By reducing the candidate window by 1/6 each cycle, we were able to calculate identical thresholds to the “accurate” method in seconds rather than hours. This opens up the ability to perform efficient Costes Colocalization calculations on 16-bit images (Figure 5B). In theory these accelerated methods could ‘overshoot’ the target threshold by a small margin in rare instances, but in our testing they consistently produced identical results to the “accurate” implementation. Nonetheless we have made all three strategies (“accurate”, “fast” and “faster”) available within the module settings. Other colocalization methods did not suffer from the same degree of performance issues, but additionally updating them to newer implementations reduced the time taken for the module to process without Costes features enabled (Figure 4C).

**Figure 5.**
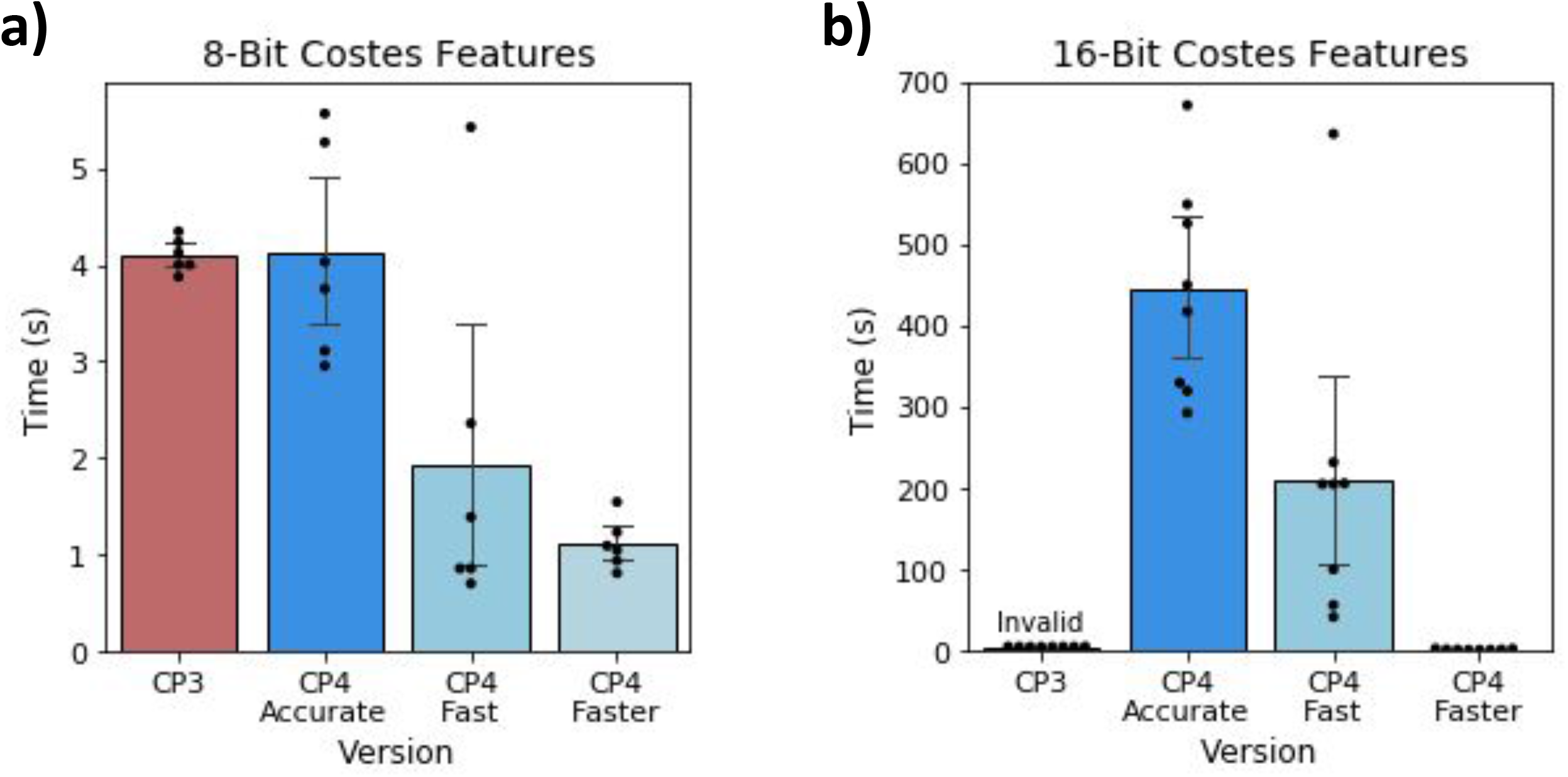
Performance of alternative Costes automated thresholding strategies. Execution times for the MeasureColocalization module performing 1 pairwise comparison with Costes features enabled, using each algorithm on (A) 8-bit images from the ExampleFly pipeline (n=6) or (B) 16-bit images from the example Cell Painting dataset (n=8). On 16-bit images results from CellProfiler 3 are calculated incorrectly, but shown to illustrate relative performance.

Together, these improvements will substantially reduce the computational time and power necessary to process images, particularly when working with large, complex data sets. This will have the added benefits of reducing resource costs for researchers, making large-scale analysis with CellProfiler more affordable and accessible. The reduction in analysis time will also reduce the environmental impact of running such pipelines.

### Performance in common workflows

To examine the impact of our changes on performance on a large heterogeneous workflow, we compared the performance of CellProfiler 3 to CellProfiler 4 when running the Cell Painting assay protocol (11). This workflow is typically performed on large datasets in a cluster environment, so we selected a sample of 48 image sets from a published dataset and measured processing times on a single machine. Execution times were captured for each module across three independent runs of this dataset. The sum of these timings represents the total workload executed by each module, excluding file I/O operations. These measurements revealed a 10-fold reduction in total CPU time required to analyze each image (Figure 6A).

**Figure 6.**
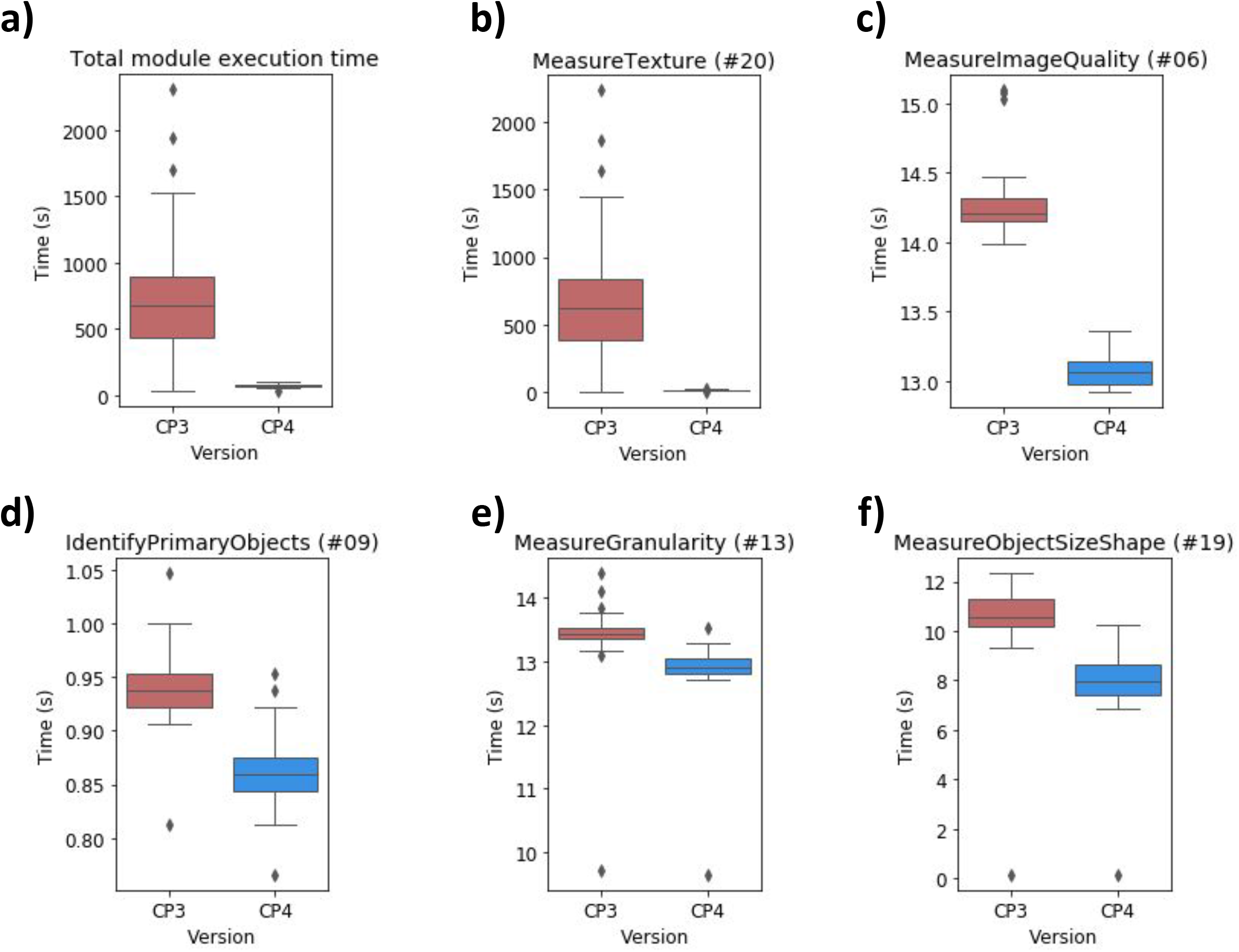
Performance of selected modules within the Cell Painting assay protocol. Numbers in brackets within panel titles correspond to modules in Figure S1. (A) Total module execution time (measured in CPU time) per image set for all modules in the pipeline. (B) Execution time for the MeasureTexture module per image set. (C) Execution time for the MeasureImageQuality module per image set. (D) Execution time for the IdentifyPrimaryObjects module per image set. (E) Execution time for the MeasureGranularity module per image set. (F) Execution time for the MeasureObjectSizeShape module per image set.

In keeping with our expectations, the refinements to MeasureTexture contributed the majority of the performance improvements that we observed (Figure 6B). We also noted small improvements in the MeasureImageQuality (Figure 6C), IdentifyPrimaryObjects (Figure 6D), MeasureGranularity (Figure 6E) and MeasureObjectSizeShape (Figure 6F) modules. Other modules in the pipeline exhibited similar performance in both versions or took negligible time to execute (Figure S1).

## Discussion

As the adoption of high content microscopy methods continues to expand there may be several areas where CellProfiler could be expanded with new functionality. Analysis of tissue sections stands out as a potential area of improvement. The large file sizes associated with tissue specimens pose a challenge for image analysis, as system memory typically is not sufficient to load the entire image at once, a bottleneck which could be avoided by adopting packages such as Dask (13) as a means of handling such images by loading subsections of an image on-demand. This would expand the utility of CellProfiler within the digital pathology field.

We also aim to continue adding support for 3D analysis to modules that currently only support 2D workflows. While segmentation is possible in 3D pipelines, additional tools and measurements will be valuable for laboratories using CellProfiler. Alongside this, further performance improvements will continue to benefit researchers, particularly when working with large datasets.

The splitting of cellprofiler-core into a standalone package has also laid the groundwork for producing a stable API for use in other Python-based applications. This will eventually allow users to modify and execute pipelines from within environments such as Jupyter, which may be of benefit to researchers looking to automate complex workflows. This API could provide a higher-level interface for common image processing tasks such as object segmentation, which would simplify the workflow for analyzing images directly within a Python environment and could serve as a bridge to Python tools whose GUI is incompatible with CellProfiler’s, such as Napari (14). The current implementation provides access to all of CellProfiler’s important classes and the ability to run pipelines or individual modules. For future development we would like to introduce a more convenient system for programmatically generating image sets without the need for the original input modules or CSV files.

In recent years there has been considerable development towards deep learning models which can perform image segmentation in an automatic manner. Providing access to these algorithms would be of substantial benefit to CellProfiler’s users, however the need for dedicated hardware and software to run these models poses a challenge for packaging and distribution. To avoid compatibility issues with older hardware, as well as to minimise the software dependencies needed to run CellProfiler, our approach has been to develop independent plugin modules which are distributed separately from the main CellProfiler program. For CellProfiler 3 we previously released a plugin for NucleAIzer (15), and in the future we hope to investigate integrations with other popular models such as Cellpose (16) and Stardist (17).

## Conclusions

The migration of CellProfiler to Python 3 will ensure that the software will remain accessible and maintainable in the coming years. In CellProfiler 4 we have further refined the user interface and introduced new modules and features to help scientists to develop and execute their analysis workflows. The targeted performance improvements in this version will substantially reduce computational costs associated with high throughput image analysis, broadening the potential applications for this open-source software package.

## Availability and Requirements

Project name: CellProfiler

Project home page: https://cellprofiler.org/

Operating system(s): Windows, MacOS, Linux

Programming language: Python 3

Other requirements: Java 1.6+ (JDK 14 bundled with builds)

License: BSD 3-Clause License

Any restrictions to use by non-academics: None

## List of Abbreviations

API: Application Programming Interface
CP3: CellProfiler 3
CP4: CellProfiler 4
I/O: input/output

## Declarations

### Ethics approval and consent to participate

Not applicable.

### Consent to publish

Not applicable.

### Availability of data and materials

CellProfiler 4 is open-source software which has been made freely available to the scientific community. Pre-compiled builds for Windows and MacOS, as well as documentation manuals, are available at http://cellprofiler.org. Source code is available at https://github.com/CellProfiler/CellProfiler. Benchmarking and visualizations presented in Figures 1C, 1D, 1F, 3A-C, 4B-C, and 5A was performed with the publicly available pipelines and image set “ExampleFly”(https://github.com/CellProfiler/examples/tree/master/ExampleFly)). Benchmarking and visualizations presented in Figures 1A, 3D, and 4A were performed with the publicly available “3D Monolayer” pipeline and image set which can be accessed at https://github.com/CellProfiler/tutorials/tree/master/3d_monolayer. Cell Painting benchmarking experiments in Figures 5B, 6, and S1 made use of a previously published data set (Plate 37983 from https://bbbc.broadinstitute.org/BBBC025) and pipeline (analysis.cppipe from https://github.com/carpenterlab/2016_bray_natprot/blob/master/supplementary_files/cell_painting_pipelines.zip, cited in (11)). The pipeline for this data set was originally written for CellProfiler 2, and so was adjusted to run on CellProfiler 3 and CellProfiler 4 with comparable outputs. These adjusted pipelines as well as the sample data and pipeline used to produce Figure 2 are provided in a public GitHub repository (https://github.com/carpenterlab/2021_Stirling_submitted).

### Authors’ contributions

DRS, MJS-B, AML, BAC, and AG developed the software. DRS wrote the manuscript, with editorial contribution and supervision from AEC and BAC.

## Acknowledgements

The authors would like to thank Katrin Leinweber, Vito Zanotelli, Erin Weisbart, Chris Allen, Nasim Jamali, and Pearl Ryder for contributions to bug fixes and testing of pre-release versions of this software. We also thank all the members of the bioimaging community who have provided feedback and suggestions which have helped to guide this work.

---

**Figure S1.**
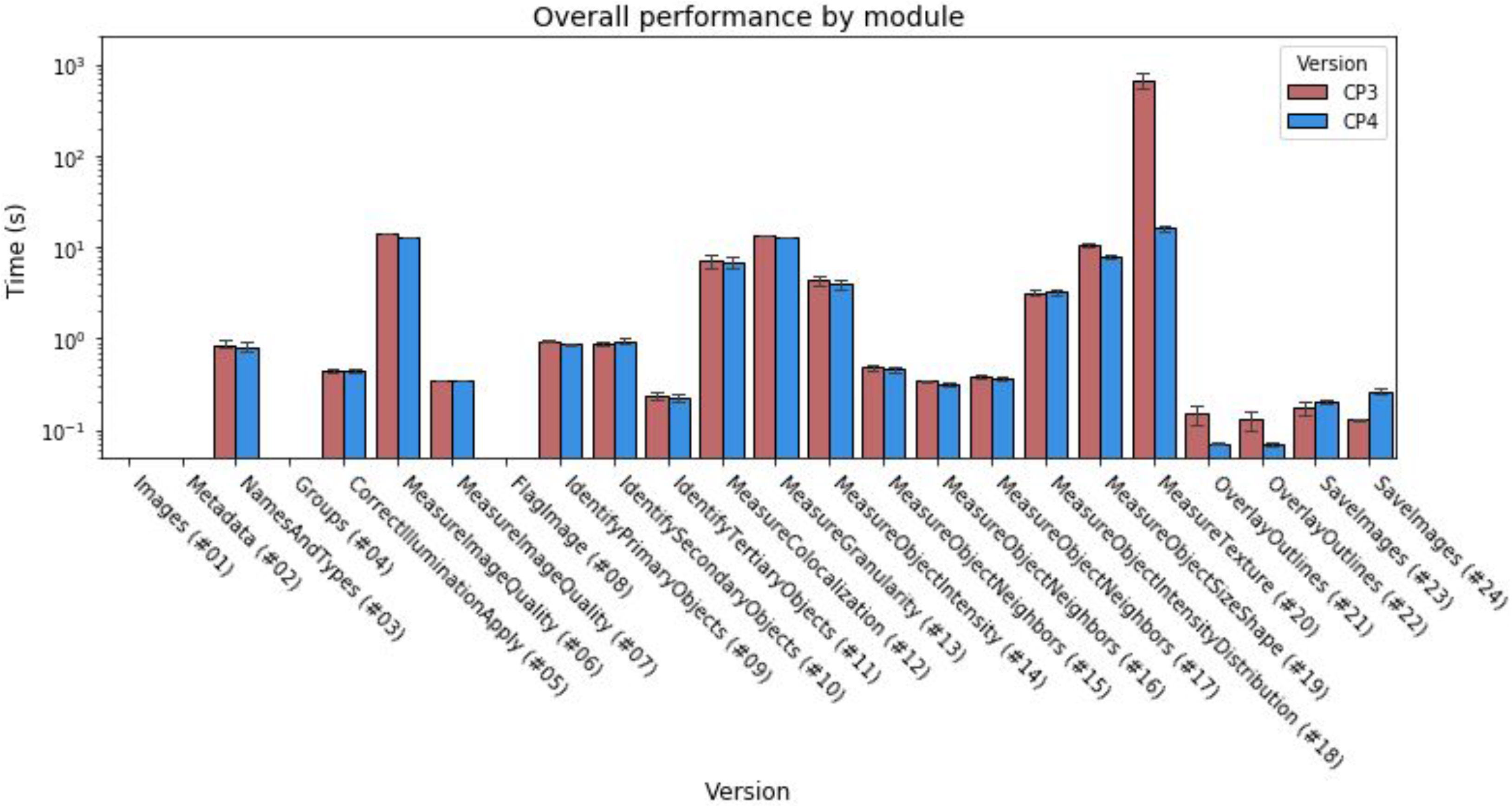
Execution times of all modules within the Cell Painting example pipeline. Measured as per-image CPU time taken for each module in the Cell Painting assay protocol (n=48). I/O loading operations in the Images module are not recorded by these measurements.

